# Cohesin dependent compaction of mitotic chromosomes

**DOI:** 10.1101/094946

**Authors:** Stephanie A. Schalbetter, Anton Goloborodko, Geoffrey Fudenberg, Jon M. Belton, Catrina Miles, Miao Yu, Job Dekker, Leonid Mirny, Jon Baxter

## Abstract

Structural Maintenance of Chromosomes (SMC) protein complexes are key determinants of chromosome conformation. Using Hi-C and polymer modelling, we study how cohesin and condensin, two deeply-conserved SMC complexes, organize chromosomes in budding yeast. The canonical role of cohesins is to co-align sister chromatids whilst condensins generally compact mitotic chromosomes. We find strikingly different roles in budding yeast mitosis. First, cohesin is responsible for compacting mitotic chromosomes arms, independent of and in addition to its role in sister-chromatid cohesion. Cohesin dependent mitotic chromosome compaction can be fully accounted for through *cis*-looping of chromatin by loop extrusion. Second, condensin is dispensable for compaction along chromosomal arms and instead plays a specialized role, structuring rDNA and peri-centromeric regions. Our results argue that the conserved mechanism of SMC complexes is to form chromatin loops and that SMC-dependent looping is readily deployed in a range of contexts to functionally organize chromosomes.

**Highlights:**

- Cohesin compacts mitotic chromosomes independently of sister chromatid cohesion.
- Formation of *cis*-loops by loop extrusion fully accounts for cohesin-mediated compaction.
- Condensin is not required for mitotic chromosome compaction of yeast chromosome arms
- Condensin has a focused pre-anaphase role at centromeres and rDNA in yeast

## Introduction

The extreme length of chromosomal DNA requires organizing mechanisms to promote faithful chromosome segregation when cells divide. Microscopy and 5C and Hi-C studies indicate that mitotic chromosomes are compacted by the introduction of an array of intra-chromosomal (*cis*) DNA loops across the chromosome (Dekker and Mirny, 2016; Earnshaw and Laemmli, 1983; Liang et al., 2015; Naumova et al., 2013). Such looping of chromosomes is predicted to compact chromosomes *in cis* whilst resolving different chromosomes *in trans*. The mechanism by which mitotic intra-chromosomal loops are generated is unresolved.

The structural maintenance of chromosome (SMC) complexes are universally required for normal mitotic chromosome compaction and faithful chromosome segregation, and are key candidates for the *cis*-looping function (Fudenberg et al., 2016; Gruber, 2014; Hirano, 2016). SMC complexes are remarkably conserved from bacteria to higher eukaryotes and share a common complex architecture (Uhlmann, 2016).

Despite their common architecture and enzymology, SMC complexes appear to have distinct roles in chromosome organization through the cell cycle. In metazoans, the SMC condensin complex is required for full chromosome compaction in extracts and in cells (Hirano, 2016; Hudson et al., 2009). *In vitro* and *in vivo* studies suggest that condensin functions by forming *cis*-loops and localizing to the chromosomal axes (Hirano, 2012; Paulson and Laemmli, 1977). In contrast, the SMC cohesin complex is traditionally thought to maintain cohesion between the sister chromatids, from G2 until the metaphase to anaphase transition (Uhlmann, 2016). Intriguingly, mounting evidence suggests that cohesin may also act in interphase to stabilize distal chromosomal interactions (Dowen et al., 2014; Hadjur et al., 2009; Kagey et al., 2010).

How these distinct *in vivo* functions are produced by complexes with apparently similar enzymatic capabilities is a topic of active debate. Topological entrapment (Cuylen et al., 2011; Haering et al., 2008; Wilhelm et al., 2015) of replicated chromosomes in *trans* by the cohesin complex is thought to promote sister-chromatid cohesion (Uhlmann, 2016). A similar mode of entrapment of distal regions of DNA in *cis* by condensin has also been proposed to generate DNA looping in mitosis by stabilizing random distal DNA-DNA interactions (Cheng et al., 2015). Although random cross-linking of chromosomes by protein bridging could generate loops, such looping would not be able to differentiate between *cis* and *trans* interactions (Alipour and Marko, 2012) and would disrupt rather than promote chromosome resolution and segregation (Goloborodko et al., 2016b). To overcome such issues an alternative mechanism of *cis*-loop formation has been proposed, called DNA loop extrusion (Alipour and Marko, 2012; Nasmyth, 2001; Riggs, 1990). A DNA loop-extruding enzyme binds to a region of DNA and then spools in adjacent DNA until the enzyme either unbinds or its progression is blocked (Fudenberg et al., 2016; Goloborodko et al., 2016b; Sanborn et al., 2015). *In silico* simulations demonstrate that loop extrusion can compact and segregate chromosomes in prophase, and form domains during interphase, suggesting that condensin and cohesin may be constituents of DNA loop extruding machines in mitosis and interphase, respectively (Fudenberg et al., 2016; Goloborodko et al., 2016a). Topological entrapment of DNA by SMC complexes is consistent with SMC complexes being involved in a loop extrusion mechanism.

A wealth of well-characterized mutants makes budding yeast an ideal model system for the study of the fundamental mechanisms of SMC action. However, unlike in metazoan mitosis where individual chromosomes condense into cytologically resolvable structures, mitotic chromosome compaction is optically fairly subtle in budding yeast. Indeed, apart from the bulky rDNA locus, mitotic chromosomes are not readily resolvable (Guacci et al., 1994). Nevertheless, the small genome size of budding yeast makes it an ideal candidate for Hi-C approaches, as high-quality, high-resolution datasets can be obtained at lower cost (Zimmer and Fabre, 2011). The small genome size and defined geometry of the budding yeast nucleus also makes its chromatin organization an ideal candidate for computational modeling (Duan et al., 2010; Wong et al., 2012).

Here we examine the connection between SMC complex activity, *cis*-looping and mitotic chromosome compaction in budding yeast using a combination of budding yeast genetics, Hi-C and *in silico* modeling. We find that budding yeast mitotic chromosome compaction can be fully accounted for by *cis*-looping by loop extrusion along chromosome arms. Surprisingly, mitotic looping is not dependent on condensin activity. Rather mitotic cohesin activity is required for *cis*-looping and mitotic chromosome compaction in budding yeast.

## Results

### Hi-C analysis of budding yeast mitotic chromosome compaction

To define mitotic chromosome compaction of budding yeast cells at high resolution, we used Hi-C on synchronized populations arrested in G1 or in mitosis (M) (Figure 1A and Figure S1A). The synchronized populations were fixed with formaldehyde and Hi-C libraries prepared to assess chromatin conformation in each condition (Figure 1A). Massive parallel sequencing produced on average 60 million unique valid pairs for each library (Supplemental Table 1). Valid contacts were then binned to 10kb and iteratively corrected to allow direct comparison of the Hi-C contact maps from the different conditions (Imakaev et al., 2012).

**Figure 1.**
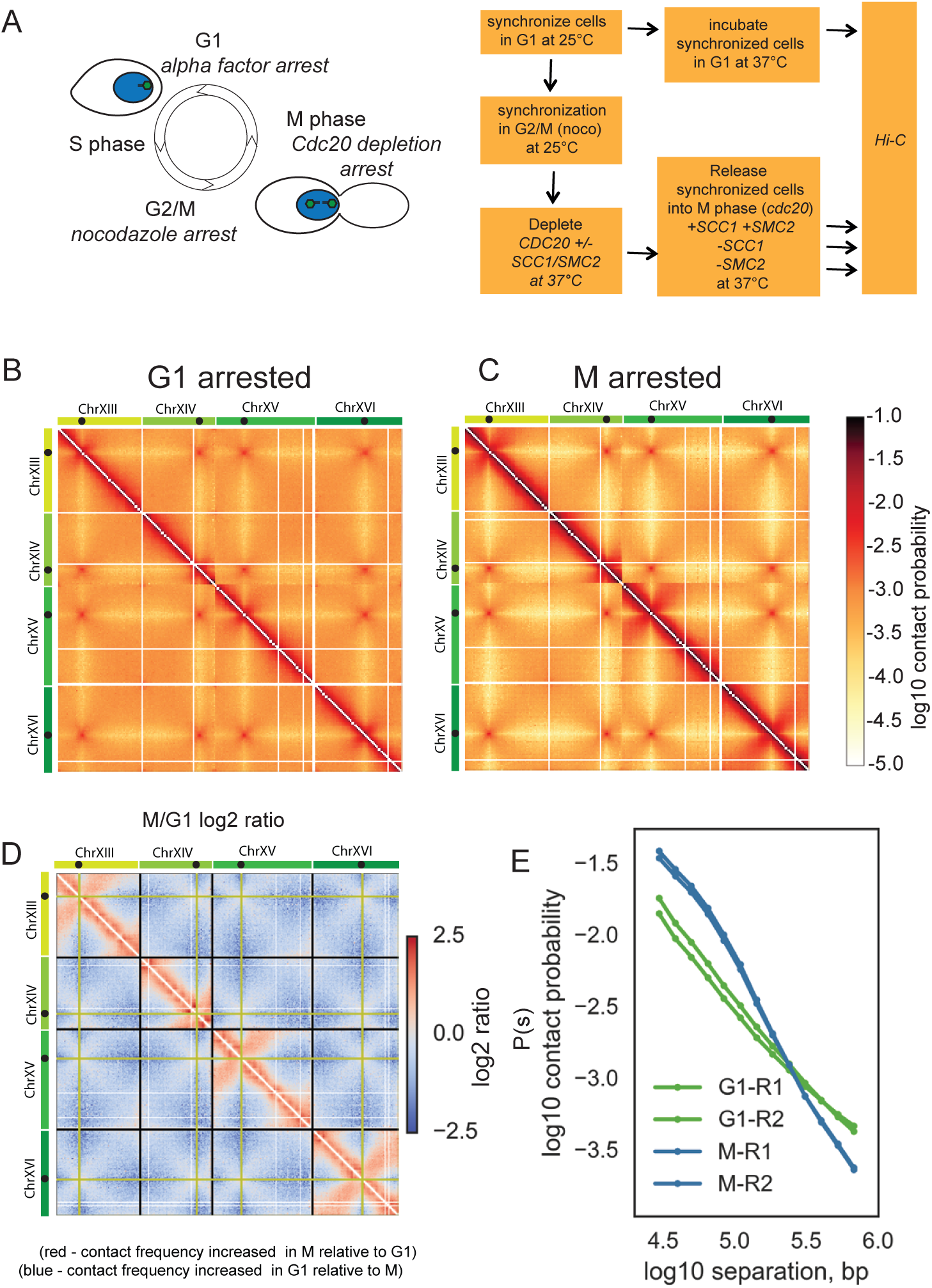
Budding yeast chromosomes are compacted in mitosis. **(A)** Experimental procedure to synchronize cells in either G1 or M, with and without cohesin and condensin. Green spots in cartoon on the yeast nucleus (blue) represent spindle pole bodies. **(B)** Hi-C contact heatmap from G1 cells synchronously arrested at 37°C with alpha factor **(C)** Hi-C contact heatmap from arrested in M phase by depletion of Cdc20 at 37°C. Both Hi-C maps have been normalized by iterative correction at 10kb resolution. Heatmap colour scale represents log10 number of normalized contacts. The Hi-C contact map for chromosomes XIII to XVI is shown as representative of the whole genome. **(D)** Log2 (M/G1) ratio of the data displayed in B and C. Regions where contact frequency was higher in M than G1 are shown in red, regions where contact frequency was lower in M than G1 in blue. **(E)** Contact probability, P(s), as a function of genomic separation, s, for G1 and M phase cells averaged over all chromosomes.

Analysis of the Hi-C contact maps in both G1 and M confirmed the presence of the main features of budding yeast nuclear organization reported previously (Duan et al., 2010); a Rabl-type organization with strong centromere clustering and arm length-dependent telomere interactions (Figure 1B, C and Figure S1B). However, comparison of the G1 and M phase contact maps (Figure 1B, C) and inspection of the log2 ratio of the two maps (Figure 1D) demonstrated that chromosomal arms became resolved from one another in mitosis relative to their interphase state. Despite centromere clustering being intensified in M, the frequency of inter-chromosomal contacts along chromosome arms was reduced compared to G1.

Concurrently, changes in local conformation were observed along chromosome arms. In M cells, the frequency of intra-chromosomal contacts less than 100kb apart were markedly increased relative to G1 whilst longer-range intra-chromosomal contacts became less frequent (Figure 1D). Analysis of intra-arm contact probability, *P(s)*, with chromosomal distance *s* (Figure 1E) demonstrated that G1 decayed at a similar rate at all distances, while M had a markedly slower decay at short distances (<100kb), suggesting chromosome compaction at this scale, followed by a more rapid decay afterwards. Analysis of P(s) of each individual chromosomal arm confirmed that these changes occurred uniformly across all chromosomes (Figure S1C). Interestingly, the two regimes of P(s) that we observed in budding yeast M, are reminiscent of Hi-C from mammalian mitotic cells which also displayed a slow decay in contact frequency followed by a more rapid drop-off (Naumova et al., 2013). Therefore, Hi-C analysis demonstrates that budding yeast chromosomes become compacted in mitosis.

### Modeling of mitotic chromosomes predicts cis-looping

We next developed polymer models to test what changes to chromosomal structure can underlie the observed changes in the G1 and M Hi-C maps. Following previous simulations (Tjong et al., 2012; Wong et al., 2012) of yeast interphase organization, we modeled the Rabl organization of the yeast genome as 16 long polymers confined to a spherical nucleus (Figure 2A-E, **Methods**). Chromosomes are tethered by the centromeres to the spindle pole body, telomeres are held to the nuclear periphery, and the whole genome is excluded from the nucleolus, located opposite the spindle pole body (Figure 2A). Following previous analysis of 3C and imaging data (Dekker, 2008) we modeled the yeast chromatin fiber as a polymer of 20nm monomers (Figure 2C), each monomer representing 640bp (~4 nucleosomes), with excluded volume interactions and without topological constraints, subject to Langevin dynamics in OpenMM (Eastman and Pande, 2010; Eastman et al., 2013). We then additionally introduced intra-chromosomal *(cis-)* loops generated by loop extrusion of varying number and coverage, i.e. the fraction of the genome spanned by all loops combined (Figure 2B), motivated by previous models of mammalian mitotic chromosomes compacted by arrays of consecutive loops (Goloborodko et al., 2016b; Naumova et al., 2013), as well as models of interphase chromosomes with extruded loops (Fudenberg et al., 2016). Since changes occurred relatively uniformly along chromosomal arms in M Hi-C maps, we introduced cis-loops stochastically from cell-to-cell at sequence-independent positions along the chromosomal arms. For each combination of loop coverage and number, we collected conformations, generated simulated Hi-C maps, and calculated simulated P(s) curves (Figure 2D, E).

**Figure 2.**
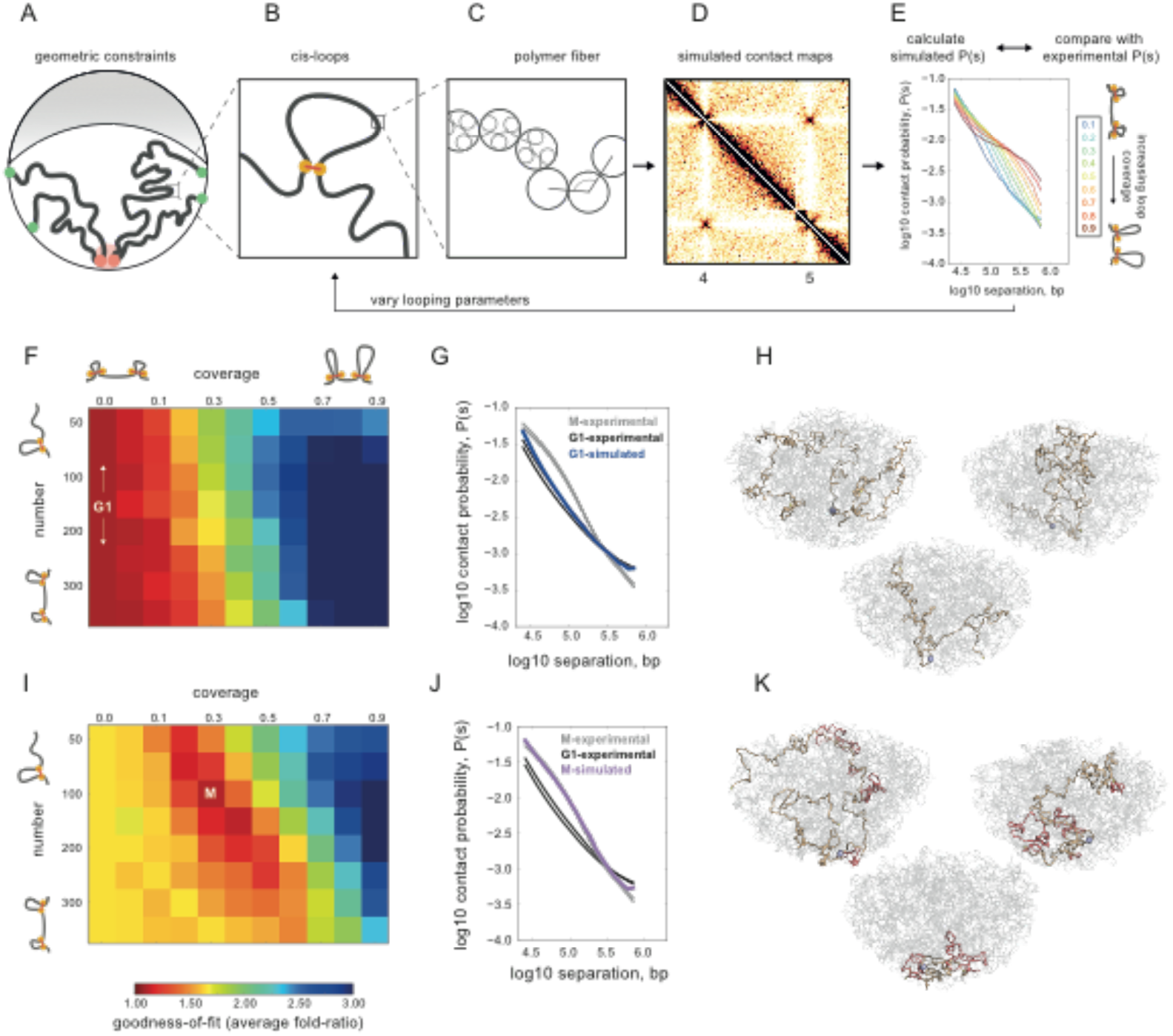
Polymer simulations of the yeast genome support compaction by intra-chromosomal loops in mitosis. **(A)** Illustration of geometric constraints used in simulations: confinement to a spherical nucleus, clustering of centromeres (red), localization of telomeres to the nuclear periphery (green), and exclusion of chromatin from the nucleolus (grey crescent). **(B)** Intra-chromosomal *(cis-)* loops, generated with a specified coverage and number per yeast genome. **(C)** Chromatin fiber, simulated as a flexible polymer. **(D)** Simulated contact maps are generated in simulations with the above constraints. **(E)** P(s) curves are then calculated from simulated contact maps. Simulations are run with systematically varied *cis-*loop parameters (coverage and number of loops), and the resulting P(s) curves are compared with experimental data. Shown here are of P(s) curves for 150 loops, and a range of coverage. **(F)** Goodness-of-fit for simulated versus experimental intra-arm P(s) in G1. Goodness-of-fit represents the average fold deviation between simulated and experimental P(s) curves, best-fitting values indicated with white text. The coverage=0.0 column represents the fit for simulations without intra-chromosomal loops. **(G)** P(s) for best-fitting G1 simulations (coverage=0.0, i.e. no-loops) versus P(s) for each experimental replica of G1 and M. **(H)** Three sample conformations from the no-loops simulations; one chromosome highlighted in light brown (clockwise from upper left: XI, V, III), with its centromere in blue, telomeres in yellow, and the rest of the genome in grey. **(I)** as F, but for experimental M Hi-C. **(J)** Best-fitting simulated P(s) for M has N=100-150, coverage=0.3-0.4. **(K)** Conformations for (N=100, coverage=0.3) with *cis*-loops additionally highlighted in light red.

Comparison of simulated and experimental P(s) curves allowed us to identify changes in chromosome organization upon transition from G1 into M-phase (Figure 2E). In G1 best fits of experimental Hi-C data were obtained when chromosomes had no *cis*-loops (coverage=0.0, Figure 2F-H,). In contrast, in mitosis we found that *in silico* models with ~10 loops per Mb, ~35kb each, covering ~35% of the genome, closely reproduced the main features of the experimental Hi-C data (Figure 2I-K, and Figure S2A, with the different polymer organization of G1 and M simulated in Figure 2H and 2K). Interestingly, introducing sister chromatids to the best-fitting G1 models either with or without sister chromatid cohesion between cognate loci, could not account for the differences we observe between G1 and M chromosomes (Figure S2B). Therefore, the introduction of cis-loops into budding yeast chromosomes by a mitosis-specific activity accounts for the differences we observe between G1 and M chromosomes.

### Cohesin is required for mitotic cis-looping, independently of cohesion

Next, we sought to identify the factors responsible for formation of these *cis*-loops in yeast mitosis. In budding yeast, *in situ* hybridization visualization of chromosomes has indicated that both cohesin and condensin are required for chromosome condensation (Freeman et al., 2000; Guacci et al., 1997). Cohesin is required for pre-anaphase rDNA chromosome condensation whereas condensin is required for both pre- and post-rDNA condensation. Also both the SMC complexes are inactive in G1 and active in M (Hu et al., 2015; Lavoie et al., 2004; Robellet et al., 2015; Uhlmann et al., 1999). As either could be responsible for imposing loops in yeast mitosis, we genetically ablated cohesin or condensin activity in mitotically arrested cells using defined inducible mutations and assayed the changes in chromosomal structure by Hi-C.

We first examined the role of cohesin in budding yeast mitotic chromosome condensation using the *scc1-73 ts* allele of cohesin. Under the restrictive conditions *scc1-73* (S525N) loses its affinity for Smc1/3 resulting in loss of cohesin complex function (Haering et al., 2004; Michaelis et al., 1997). We pre-synchronized cells in G2/M under the permissive conditions, before incubating the cells at the restrictive temperature (Figure 1A). We then examined the mitotic chromatin conformation of cells in the absence of cohesin.

Loss of mitotic cohesin activity led to a disappearance of the characteristic mitotic features, as determined from Hi-C (Figure 3A-C), despite being maintained in metaphase by the depletion of Cdc20 (Figure S1A). Indeed, the two-regime M-phase P(s) disappeared, becoming closer to that of G1 (Figure 3D), with diminished short distance (<100kb) contacts and more frequent longer-range and inter-chromosomal contacts (Figure 3A-D). These changes were not only observed along chromosome arms connected to centromeres. Cohesin depletion also resulted in loss of short distance (<100kb) contacts and more frequent longer-range and inter-chromosomal contacts in the post-rDNA regions of chromosome XII (Figure S3). This region is isolated from the centromere by the rDNA array and not subject to any potential indirect effects resulting from cohesin action at the centromere. Therefore, cohesin depleted mitotic chromosomes lose mitotic compaction. Consistently, modeling indicated that Hi-C maps of cohesin mutants were well-fit by simulations with many fewer loops than wild-type mitotic Hi-C maps (Figure 3E). Together our results indicate that cohesins are required for mitotic *cis*-loops along chromosome arms.

**Figure 3.**
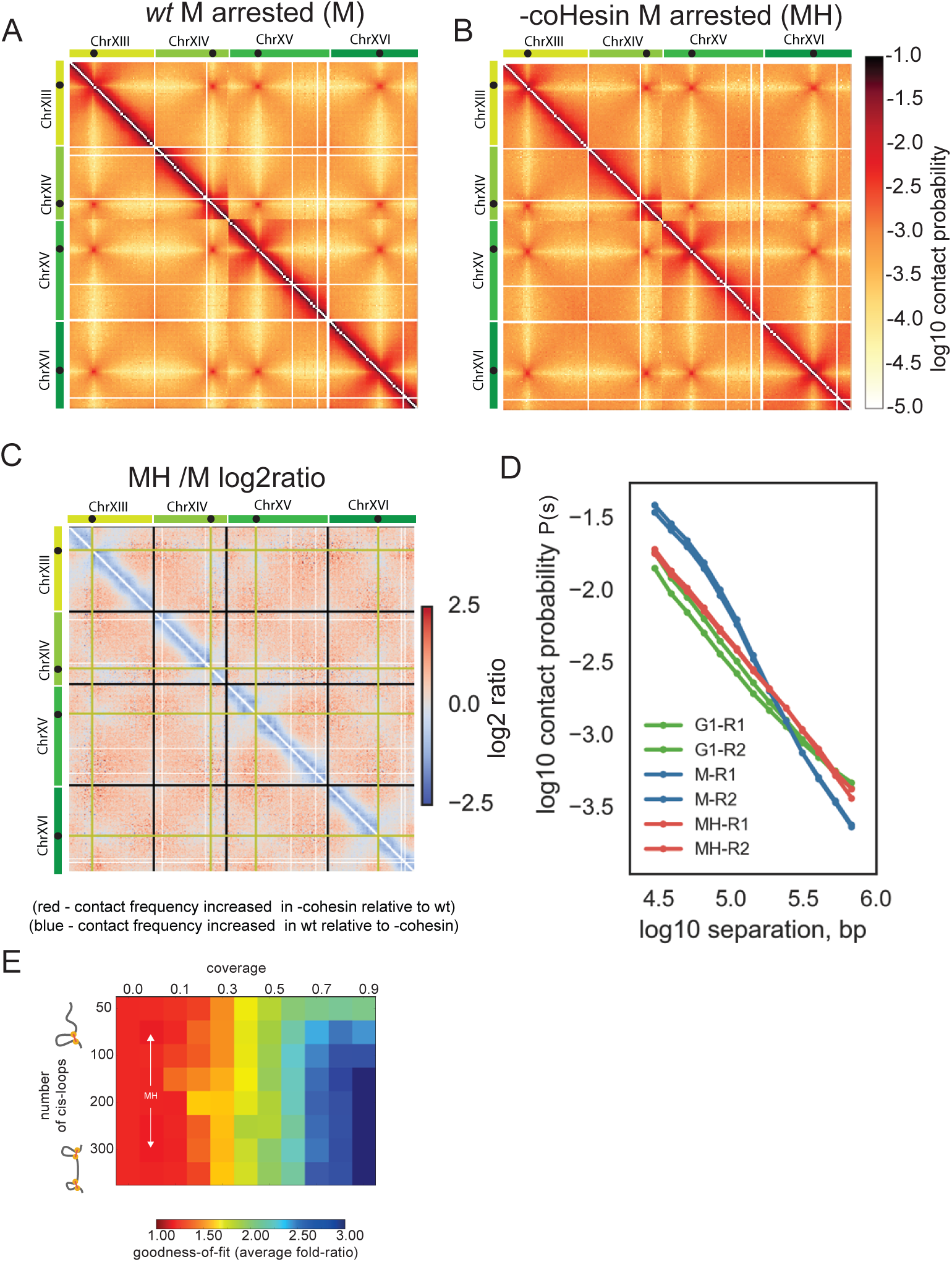
Cohesin activity is required for mitotic compaction and cis-looping. **(A)** Hi-C data collected from M phase (reproduced from Figure 1C) **(B)** Hi-C data collected from M phase cells following disruption of coHesin using the *scc1-73* allele (MH). Chromosomes XIII to XVI are shown as representative of the whole genome. **(C)** Log2 ratio of *–cohesin* MH dataset over wt M dataset (MH/M), displayed in B and A, respectively. **(D)** Contact probability (P(s)) for M and MH phase cells and G1 cells averaged over all chromosomes. **(E)** Goodness-of-fit for models with variable *cis*-loop coverage (horizontal axis) and number (vertical axis).

To further test for a cis-looping role of cohesin in mitosis, we assessed whether cohesin could still compact mitotic chromosomes when no sister chromatid cohesion was present. We examined cells with a *cdc45* degron allele (Figure 4A), blocked in mitosis. These cells generate mitotic chromosomes without a preceding round of DNA replication and therefore without sister chromatids (Tercero et al., 2000). Confirming our hypothesis regarding a mitotic *cis*-looping role of cohesin, chromosomes had contact frequencies in mitosis distinct from G1, exhibiting P(s) with two regimes, similar to that in wt M-phase Hi-C, and indicative of the presence of *cis*-loops (Figure 4B). This feature disappeared upon cohesin depletion, even without the presence of a sister chromatid (Figure 4B, C). Moreover, loss of cohesin function also led to loss of chromosome resolution with increased inter-arm interactions (Figure 4C). Together with our *in-silico* analysis that sister-chromatid cohesion alone could not account for mitotic contact maps, our results strongly suggest a mitotic *cis*-loop forming function of cohesin, independent of, and in addition to, its accepted role in sister-chromatid cohesion.

**Figure 4.**
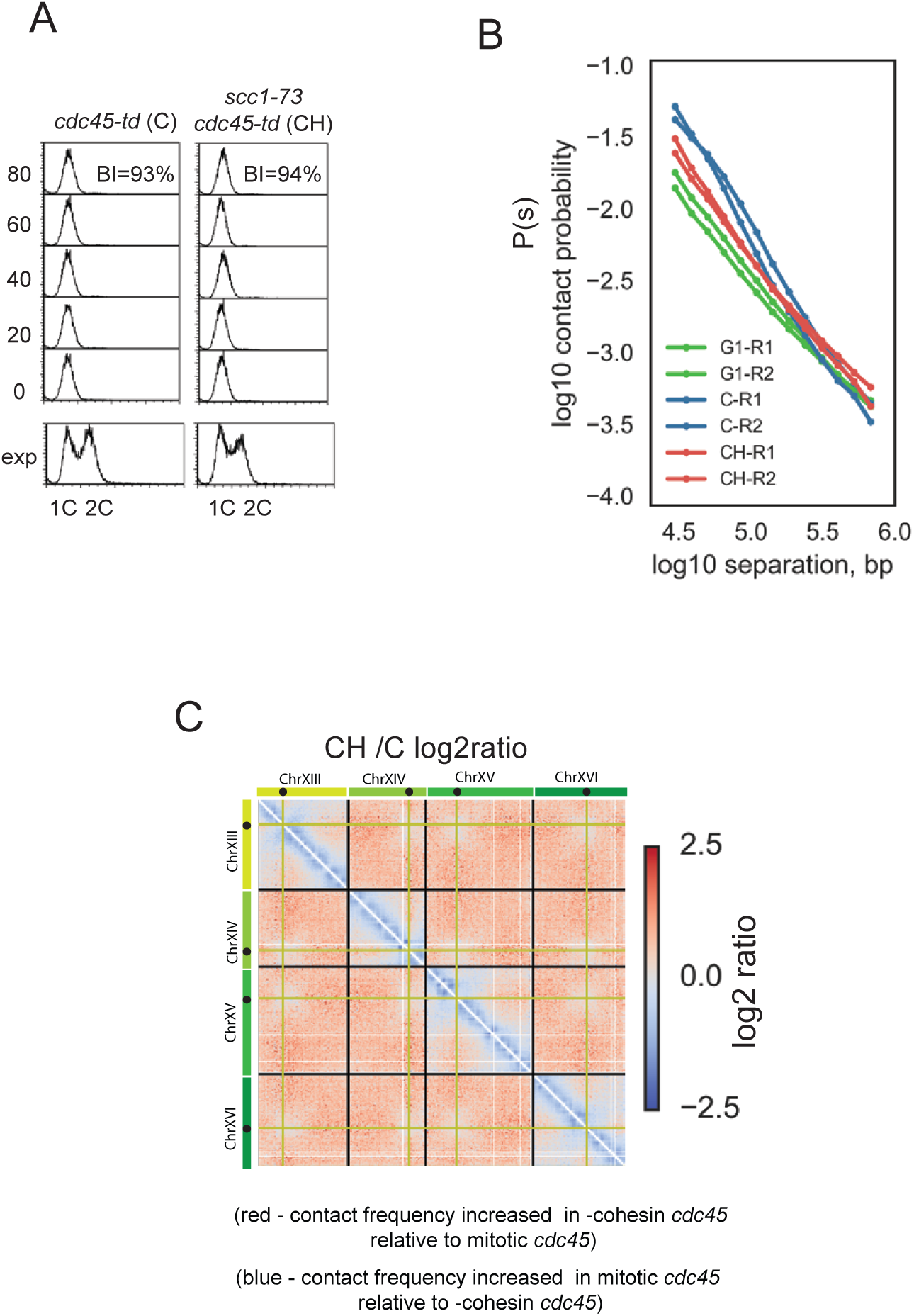
Mitotic cohesin dependent conformational changes are independent of sister chromatid cohesion. **(A)** FACS analysis of DNA content and budding analysis of *cdc45-td* (C) and *cdc45-td scc1-73* (CH) cells following release from G1 arrest into a nocodazole enforced mitotic block. Budding index (BI) confirmed that mitotic cells had activated CDK while FACS of DNA stained cells confirmed no DNA replication has taken place. **(B)** Contact probability, P(s) versus genomic separation, s, for Hi-C of mitotic *cdc45-td* (C) mitotic *cdc45-td scc1-73* (CH), and wt G1 cells (G1). **(C)** Log2 ratio of – *cohesin* C dataset over C dataset (CH/C),

### Condensin has a focused role in mitotic structure and is not required for cis-looping on chromosome arms

We next investigated the role of the other key evolutionarily conserved SMC in budding yeast, condensin, by Hi-C. We first examined the consequence of degrading condensin subunit Smc2 in mitosis using a degron allele of *SMC2*. We degraded Smc2 protein in G2/M before arresting the cells in M (Figure 1A). In contrast to cohesin, loss of condensin activity had surprisingly mild effects on mitotic intra-arm chromatin organization (Figure S4A).

Given the mild phenotype of the condensin degron, we considered the possibility that we were not completely ablating condensin function. Therefore we engineered a system predicted to cause a close-to-null condensin inactivation. We used a conditional depletion/expression system to express an enzymatically dead form of Smc2 (*SMC2K38I*) in G2/M cells whilst also depleting active degron-tagged Smc2 before arresting the cells in M. The inactive *SMC2K38I* protein is incorporated into the condensin complex and further decreases ChIP enrichment of the condensin complex on chromatin over the depletion alone (Figure S4B). Additionally, *SMC2K38I* expression increases the extent of aneuploidy that occurs following a condensin depleted mitosis (as shown by the increase in cells with <1 or >1C DNA content following cell division - Figure S4B). Therefore, we assume that expression of *SMC2K38I* prevents stable binding of condensin to chromatin and therefore approximates a null mutation for condensin activity.

We then re-examined the role of condensin by performing Hi-C on mitotic cells with our engineered null mutation for condensin activity. Despite the additional loss of condensin function generated by *SMC2K38I*, we still did not observe any loss of chromatin compaction characteristic in the arms of mitotic budding yeast chromosomes (Figure 5A-C). In contrast to cohesin depletion, the two regimes of mitotic P(s) persisted in the condensin mutant (MD) cells (Figure 5D). Consistently, simulations did not support great differences in the number of coverage of *cis*-loops along chromosomal arms (Figure 5E). We conclude that, unlike cohesin, condensin is not generally required for mitotic chromosome arm compaction in budding yeast.

**Figure 5.**
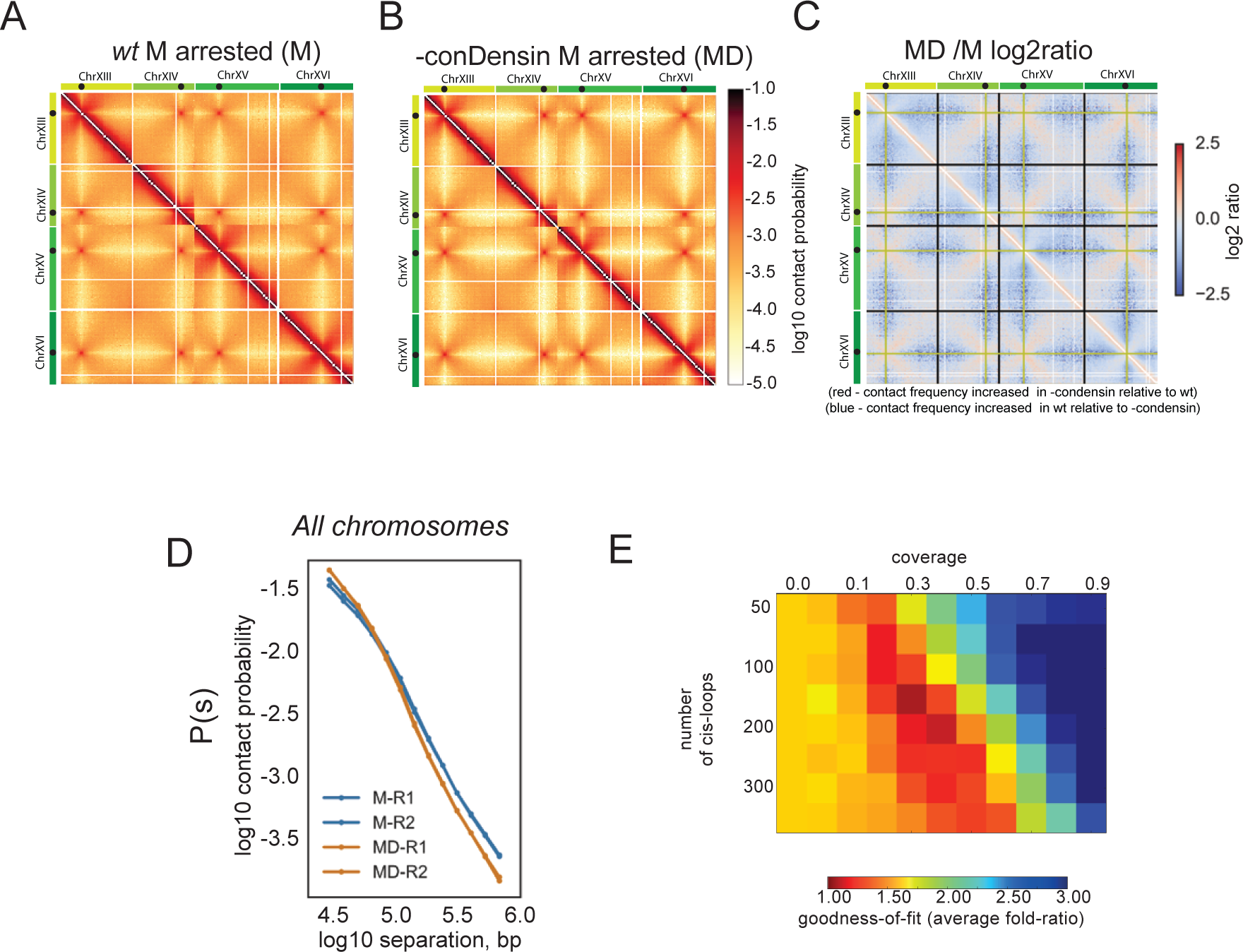
Condensin action is not required for mitotic cis-looping along chromosome arms. **(A)** Hi-C data collected from M phase cells following disruption of conDensin with *smc2td GAL1-SMC2K38I* allele (MD). Chromosomes XIII to XVI are shown as representative of the whole genome. **(B)** Log2 ratio of –condensin M dataset over wt M dataset (MD/M) **(C)** Contact probability (P(s)) for M and MD cells for all chromosomes **(D)** Goodness-of-fit for simulated versus experimental intra-arm P(s) in G1, as in Figure 2, for conDensin depleted cells.

However, visual inspection of Hi-C maps revealed that condensin activity was required for higher order chromosome structure in two specific genomic contexts, at centromeres and adjacent to the rDNA, consistent with previous studies (Freeman et al., 2000; Lavoie et al., 2002; Stephens et al., 2011). Firstly, the isolation of CEN regions from the chromosomal arms imposed by the Rabl conformation is increased in the condensin (MD) mutant, seen as the decreased contact frequency between centromeres and other regions *in cis* and *in trans* Hi-C maps (Figure 6A, B). This change is concurrent with increased co-alignment of chromosomal arms *in trans*, seen as the increased contact frequency between regions at equivalent genomic distances from the centromere (Figure 6B). Both changes are consistent with a picture where centromeres are more tightly clustered in the absence of condensin activity. Secondly, while the repeated structure of the rDNA makes it refractory to direct analysis by Hi-C, we could analyze changes in adjacent regions. Surprisingly the pre-rDNA region of ChrXII (defined as the region between CENXII and the rDNA repeats) showed significant changes in its chromosome conformation in the absence of condensin (Figure 6C-E). Interestingly, the pattern of contact changes were distinct from those lost following cohesin disruption, with contacts >100kb appearing significantly reduced (Figure 6C-E). While the pre-rDNA region displayed these distinct condensin dependent changes, the post-rDNA region, which is megabases away from the nearest centromere, remained remarkably similar to WT mitotic cells (Figure 6C, D), exhibiting the same P(s) as wildtype cells (Figure 6E). This suggests regions isolated from the centromere cluster do not incur any altered conformation in the absence of condensin. Our interpretation of the locus specific and complex conformation following condensin depletion is that condensin function is focused at centromeres and the rDNA repeats. Loss of the condensin dependent higher order structure around centromeres and the rDNA is likely to have both direct local and indirect distal effects.

**Figure 6.**
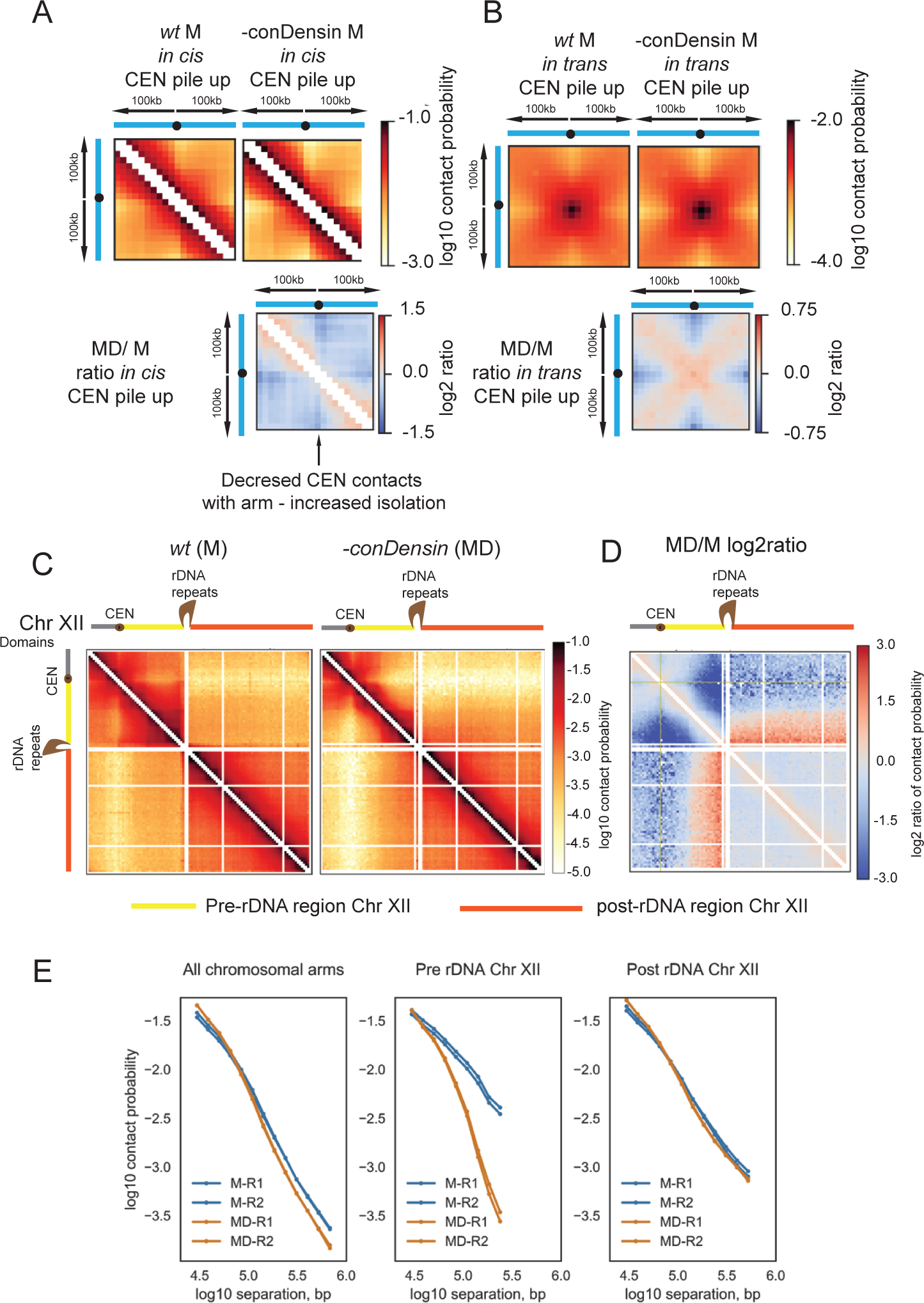
Pre-anaphase condensin activity is focused on centromeres and proximal to the rDNA repeats. **(A, B)** Pile-ups of the contact heat-maps of the 100kb peri-centromeric regions **A**) *in cis* or **B**) *in trans* on either side of budding yeast CEN sequences. Bottom, log2 ratio of the different pile-ups in the mitotically arrested state. (**C**) Hi-C contact heat maps of ChrXII in M or MD. In the cartoon representation, ChrXII is separated into three regions, the pre-CEN region (grey), the region between CEN and the rDNA repeats (yellow) and post rDNA (orange). (**D**) Log2 ratio of MD over M dataset (MD/M) for ChrXII. (**E**) Contact probability (P(s)) for M and MD cells for all chromosomes (taken from 5D) compared to contact probability (P(s)) of the pre- rDNA region and post-rDNA region of ChrXII for M and MD cells.

Finally we tested, and ruled out, the previously reported condensin-dependent tRNA gene clustering (Haeusler et al., 2008; Thadani et al., 2012). If tRNA genes were to cluster via direct bridging interactions, we would observe peaks in contact frequency between pairs of tRNA genes in the Hi-C map (Fudenberg et al., 2016). To test for such peaks with high sensitivity, we computed the average Hi-C contact frequencies between pairs of tRNA loci *in cis* and *in trans*. We did not observe any preferential contact patterns associated with tRNA pairs, either in wild-type or mutant cells (Figure S5). Thus our analyses indicate that previously reported tRNA clustering via FISH was likely produced by the global features of yeast chromosomal architecture, e.g. overall polarization resulting from a Rabl organization. Indeed, our data indicate that condensin activity can indirectly alter general nuclear organization due to direct condensin dependent effects at the nuclear organizing hubs of the centromere cluster and the rDNA array. Together our observations emphasize that in a closed system, such as a nucleus, changes at central chromosome organizing points, can lead to significant changes in relative chromosome organization in distal regions.

### Cohesin dependent loop extrusion accounts for the looping on mitotic chromosomes

Notwithstanding the complex effects of condensin action in budding yeast mitosis, our data demonstrate that cohesin dependent looping of chromosome arms is required for mitotic compaction. Two distinct mechanisms of SMC-mediated DNA looping have been proposed (Figure 7A): (i) random point-to-point loop formation stabilized by SMC complex bridging (Cheng et al., 2015); (ii) DNA loop formation by loop extrusion (Alipour and Marko, 2012; Fudenberg et al., 2016; Goloborodko et al., 2016a; 2016b). Loop extrusion can form exclusively *cis*-loops within arms but can be blocked by insulating protein complexes. Point-to-point looping relies on spatial interactions and can bridge arms that frequently contact each other across the centromere due to the Rabl orientation. The high resolution of our Hi-C data allowed us to test the different predictions of these two mechanisms. Comparison of cohesin depleted and wild-type Hi-C maps shows cohesin dependent intra-chromosomal contacts <100kb along all chromosome arms, and a noticeable lack of cohesin dependent contacts across centromeres in *cis* (Figure 7B, C, Figure S6). While the Rabl configuration enforces frequent contacts between the peri-centromeric regions on either side of the centromere, as can be seen in both wt and cohesin depleted cells (Figure 7B, C, Figure S6), only intra-arm contacts are enhanced by cohesin activity in the wild-type. This lack of cohesin dependent enhancement of cross-centromere contacts between spatially juxtaposed regions is difficult to reconcile with a bridging mechanism of loop formation. In contrast, these observations are consistent with loop extrusion activity of cohesin that is blocked by the centromere-associated multiprotein kinetochore complex. Here, kinetochores appear to act in a similar manner to DNA-bound CTCF proteins in mammalian cells that block loop extrusion by cohesin, insulating neighboring domains (Fudenberg et al., 2016 Sanborn et al., 2015). This data show that loop extrusion can fully account for the features of cohesin dependent mitotic chromosome compaction we observe.

**Figure 7.**
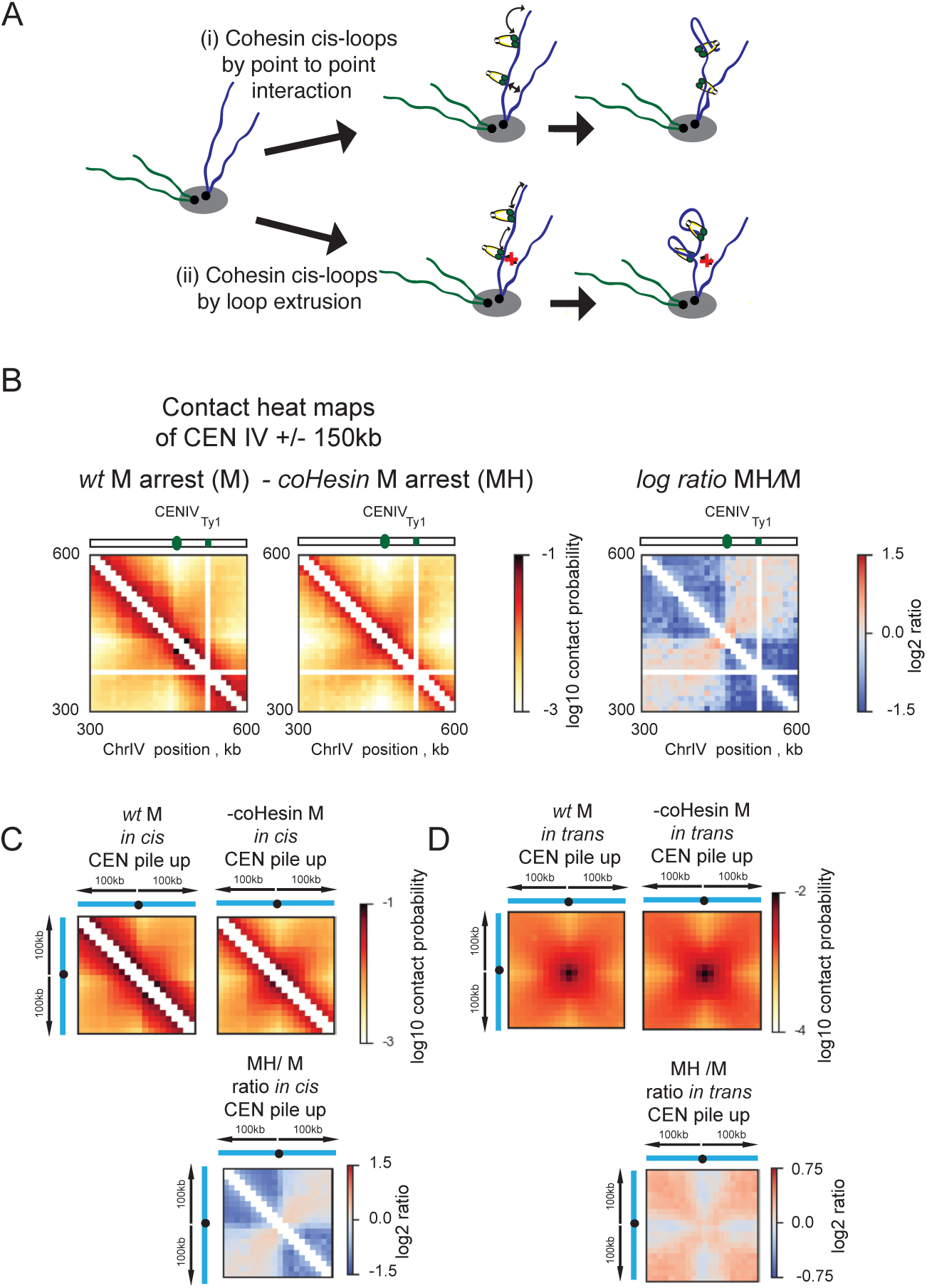
Cohesin dependent contacts around centromeres are consistent with loop extrusion activity. **(A)** Possible mechanisms of loop formation: (i) random point-to-point bridging by cohesin; (ii) loop extrusion. Since there is a high frequency of contacts between arms across the centromere due to the Rabl orientation, point to point bridging should link any DNA segments that are spatially juxtaposed and should be unaffected by distal *cis*- structures such as the centromere (top). However, a DNA tracking mechanism such as loop extrusion is likely to be blocked by a centromere, which will act analogously to a chromosomal insulator to *cis*-loop formation. **(B)** (Left) Hi-C heat maps of 150kb either side of CEN4 are shown from either mitotic cells (M) or cohesin disrupted mitotic cells (MH). (Right) Log2 ratio of –cohesin M dataset over wt M dataset (MH/M), displayed in B and A respectively. (**C**) Pile ups of interactions between pericentromeric 100kb region on either side of centromeres in *cis* of all other centromeres (excluding rDNA chromosome XII and chromosome IV) to demonstrate that centromeres act as insulators to cohesin dependent contacts between chromosomal arms, thus supporting *cis*-looping via loop extrusion. (**D**) Pile ups of interactions between pericentromeric 100kb region on either side of centromeres *in trans* between different centromeric regions demonstrates that cohesin depletion does not prevent centromere clustering (excluding rDNA chromosome XII and chromosome IV).

## Discussion

Our results support surprisingly different mitotic activities for SMC complexes of cohesin and condensin than those anticipated from their canonical functions in mammalian cells. For cohesin, our results indicate a key role in formation of mitotic intra-arm loops that are essential for chromosome compaction and resolution. For condensin, our results argue for a focused mitotic role in organizing centromeres and the rDNA locus.

The canonical role of cohesin is to establish sister-chromatid cohesion in S phase (Nasmyth, 2001). However, cohesin is still loaded onto chromosomes in M phase (Eckert et al., 2007). In addition, two populations of cohesin, one stably bound, one dynamically bound, have been found to associate with mitotic chromosomes (Chan et al., 2012; McNairn and Gerton, 2009). These findings indicate that cohesin has a function on mitotic chromosome beyond the stable juxtaposition of sister-chromatids *in trans* established in S phase. Here we show that cohesin is required for mitotic *cis*-looping, chromosome compaction and the resultant resolution of different chromosomes in budding yeast mitosis. Cohesin has also been reported to serve an interphase function in chromosome organization in higher eukaryotes and fission yeast that can be accounted for through *cis*-looping (Fudenberg et al., 2016; Mizuguchi et al., 2014; Sanborn et al., 2015). This functional coherence over long evolutionary timescales and contrasting cellular contexts argues for a fundamentally dual function of cohesin, both for the formation of DNA loops *in cis*, and holding sisters together *in trans*.

Our data indicate that cohesin is required for a loop extrusion activity in mitosis. Like other DNA tracking molecules, such as helicases, SMC complexes topologically entrap DNA, allowing rapid translocation along DNA even when chromatinized (Davidson et al., 2016; Kanke et al., 2016; Stigler et al., 2016), consistent with a direct role for SMCs in loop extrusion. Direct biochemical evidence of how SMC complexes may achieve loop extrusion and how this state is distinct from cohesive cohesin complexes is currently lacking. However, we note that the association of cohesin with chromosomes is regulated by distinct ATP hydrolysis events, which in turn are regulated by several other factors (Beckouët et al., 2016; Çamdere et al., 2015; Elbatsh et al., 2016; Murayama and Uhlmann, 2015), including known regulators of chromosome compaction (Hartman et al., 2000; Orgil et al., 2015; Tong and Skibbens, 2015). Therefore, there is clear mechanistic potential for mitotic cohesin complexes to be engaged in the distinct chromosome structuring roles of loop extrusion and sister chromatid cohesion on mitotic chromosomes.

A tempting hypothesis to account for the contrasting roles of cohesin and condensin during mitosis, is that the role of condensin has been reduced to that of an auxiliary compaction system in budding yeast. In this model condensin is deployed when compaction provided by cohesin proves insufficient or when pre-anaphase resolution of sister-chromatids is required. Indeed, the roles of condensin in providing extensive longitudinal compaction and sister-chromatid resolution are largely redundant in pre-anaphase budding yeast (Bhalla et al., 2002; Houlard et al., 2015; Ouspenski et al., 2000). In budding yeast sister-chromatids remain cohesed all across chromosome arms until anaphase (Nasmyth, 2001). Indeed, following dissolution of cohesin dependent sister chromatid cohesion few entanglements appear to remain between sister-chromatids (Farcas et al., 2011). The relatively short length of non-rDNA chromosomes also removes a general requirement for extensive longitudinal compaction for segregation. Rather, condensin dependent sister-chromatid resolution activity on budding yeast chromosome arms appears to be generally applied post-anaphase. Studies in budding yeast have shown that condensin dependent ‘adaptive hyper-condensation’ is deployed along chromosomal arms in anaphase as an emergency measure to resolve persistent entanglements (Neurohr et al., 2011; Renshaw et al., 2010). In contrast to condensin’s lack of activity on non-rDNA arms, we confirm that condensin does act prior to anaphase to mitotically re-structure two types of genomic loci likely to have exceptional resolution and segregation requirements. In budding yeast sister chromatid centromeres separate prior to anaphase (Goshima and Yanagida, 2000), consistent with a requirement for pre-anaphase condensin dependent resolution. The rDNA also has both exceptional chromosome resolution and compaction requirements for faithful segregation. The rDNA accumulates excessive levels of sister chromatid intertwines across the array (D’Ambrosio et al., 2008). In contrast to the other chromosomal arms in budding yeast, the rDNA requires extra longitudinal compaction for segregation of its extreme length (Sullivan et al., 2004). These factors could explain why centromeres and the rDNA are specifically targeted for condensin dependent action throughout mitosis. Unfortunately, the resolution of our Hi-C approach and the repetitive nature of the rDNA prevent us from analyzing the exact nature of condensin activity in re-structuring these domains. However, given the numerous connections between condensin and mitotic looping (Dekker and Mirny, 2016; Hirano, 2012) we anticipate that condensin is performing a focused loop extrusion role at these loci. In this framework, the longer and more repetitive chromosomes of higher eukaryotes, not only require functional compaction during interphase, imposed via cohesin, but will also require the additional compaction offered by condensin across all chromosomes, all the way from prophase to the end of mitosis.

Ascertaining the upper limits of cohesin dependent chromosome compaction before handover to condensin is a key question for the future. Interestingly, metazoan chromosome compaction during the mitotic cell cycle is surprisingly robust in the face of reduced condensin activity (Hudson et al., 2003; 2001), suggesting some redundancy with other chromatin looping pathways. A role of cohesin dependent looping in the early stages of higher eukaryotic mitotic compaction would be consistent with other recent studies. Increase of chromatin-associated cohesin in interphase can lead to widespread chromosome condensation (Tedeschi et al., 2014). Cohesin, like condensin, is also localized to the axis of mammalian chromosomes before its displacement in late prophase (Liang et al., 2015).

In summary our results argue that SMC complexes share an evolutionarily conserved mechanism that allows them to form chromatin loops. We speculate that the conserved mechanism of SMC action has been adapted within the different complexes to cope with the varying requirements for chromatin looping in different organisms and contexts. Unraveling how the baton of SMC function has been passed through evolution presents a fascinating topic for future research, and promises to shed light on the pleiotropic consequences of mutations to these key chromosome organizers in human disease (Watrin et al., 2016).

## Author Contributions

S. A. S. performed all cell culture and generated Hi-C libraries. A. G. analysed sequenced libraries and Hi-C data sets. G. F. modelled the conformation of the budding yeast nucleus onto the contact maps with help from A.G. J. M. B. guided Hi-C library instruction and analysed sequenced libraries. M. Y. and C. M. constructed and characterized the inducible condensin alleles. J. D. advised on study construction and guided processing and analysis of Hi-C datasets. L. M. guided the modelling of the Hi-C data. J.B. conceived and co-ordinated the study. J.B., G. F and A. G. wrote the manuscript with input from S. A. S., L.M. and J. D.

## Acknowledgements

We thank Bryan Lajoie and Johan Gibcus for aid in processing the Hi-C datasets. We thank Luis Aragon, Kim Nasmyth and John Diffley for yeast strains. This work was funded by the Biotechnology and Biological Sciences Research Council United Kingdom, (BBSRC UK) Grant ref BB/J018554/1 (S.A.S., M. Y.), the Royal Society U.K. (J. B.), NIH U54 4D Nucleome grant (DK107980) and NIH R01 (HG003143) (A. G., G. F, J. M. B., L.M., and J. D.).

